# Identification of a novel false-positive mechanism in RapidFire mass spectrometry and application in drug discovery

**DOI:** 10.1101/2025.01.24.634670

**Authors:** De Lin, Lesley-Anne Pearson, Shamshad Ahmad, Sandra O’Neill, John Post, Colin Robinson, Duncan E. Scott, Ian H. Gilbert

**Author notes:** Joint first author (De Lin, Lesley-Anne Pearson).

## Abstract

False-positives plague High Throughput Screening in general and are costly as they consume resource and time to resolve. Methods that can rapidly identify such compounds at the initial screen are therefore of great value. Advances in mass spectrometry have led to the ability to screen inhibitors in drug discovery applications by direct detection of an enzyme reaction product. The technique is free from some of the artefacts that trouble classical assays such as fluorescence interference. Its direct nature negates the need for coupling enzymes and hence is simpler with fewer opportunities for artefacts. Despite its myriad advantages, we report here a mechanism for false-positive hits which has not been reported in the literature. Further we have developed a pipeline for detecting these false-positive hits and suggest a method to mitigate against them.

**Graphical Abstract:** 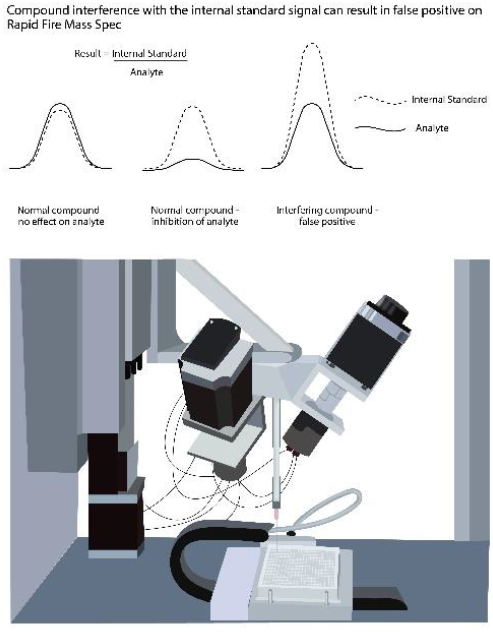

## Introduction

RapidFire mass spectrometry (RF-MS) is a high-throughput mass spectrometry technique that allows for the analysis of large numbers of samples in a short amount of time. This has made it a valuable tool for high-throughput screening (HTS) applications in various fields, such as drug discovery, metabolomics, and proteomics and is compatible with various sample matrices (cell lysate, plasma, tissue etc).^1-4^ RF-MS has been used in target-based screening for screen large libraries of compounds in various targets including *Trypanosoma brucei* AdoMetDC^5^, HIV-1 Protease^6^ and WIP1 Phosphatase^7^. The ability to assay the activity of a target by direct detection of increase of product, or depletion of substrate is a technique that is gaining in popularity, with the expectation that the removal of complex coupling reactions reduces the risk of assay interference.

An internal standard is typically added to a sample prior to mass spectrometry analysis to correct for variations in the sample preparation, instrumental parameters, and matrix effects that can affect the response of the analyte.^8^ This is particularly useful in RF-MS analysis where the sample matrices can be complex and variable, leading to variations in the analyte response.^9^ The signal ratio of the analyte to the internal standard is reported as a ‘normalised response’ for quantification. The use of internal standards can also improve the precision, accuracy, and sensitivity of RF-MS analysis. The internal standard is chosen such that it has same or similar in chemical and physical properties as the analyte but can be distinguished from the analyte by its mass-to-charge ratio (m/z) in the mass spectrometer. A stable isotope labelled (SIL) analogue is believed to be most appropriate internal standard in a quantitative mass spectrometry.^10^ However, the signal of the internal standard is potentially affected by compounds with the same molecular weight, especially in large library compound screening and when using low resolution mass spectrometry instruments (e.g. quadrupole).

We previously published a method for screening compounds in a relatively high-throughput mode against the SARS-CoV-2 protein nsp14 using RF-MS^11^ (Pearson et al 2021). We have now used this to assess the potency of a range of drug-like molecules. In addition, we used a luminescence-based MTase-Glo assay (Promega) as an orthogonal assay to confirm the activity of hits identified in the RF-MS assay. In the process of testing compounds in the RF-MS assay we identified a number of false positives with a previously undescribed mode of interference with the RF-MS. In this paper we elucidate this mode of interference as well as offering possible ways to mitigate it.

## Materials and Methods

All aqueous solutions were prepared with deionized water (Millipore, Watford, Hertfordshire, UK). All reagents were purchased from Sigma Aldrich (Gillingham, Dorset, UK) unless otherwise stated. Full-length SARS-CoV-2 nsp14 protein (DU66418) was supplied by the MRC-PPU (Dundee, UK). Nsp14 was cloned in fusion with a cleavable N-terminal GST fusion in a pGEX6P1 vector and expressed in *Escherichia coli*. Nsp14 was purified by batch purification using GSH-Sepharose beads and the tag was cleaved by PreScission protease. Cleaved nsp14 at 1.23 mg/mL was delivered in 50 mM Tris, pH 7.5, 150 mM NaCl, 270 mM sucrose, 0.1 mM EGTA, 0.03% Brij-35, 0.1% β-mercaptoethanol. Peptide IAYLKK*AT was synthesised at Cambridge Peptides by solid-phase synthesis. K^*^ refers to ^13^C and ^15^N-enriched lysine (labelled on all carbons and nitrogens).

### RapidFire assay

We have previously reported the use of RF-MS to assess the potency of compounds against the methyltransferase activity of nsp14 from SARS-CoV-2.^11^ The protein catalyses the methylation of the N7-guanosine of RNA using (S)-adenosyl methionine (SAM), which is converted to (S)-adenosyl L-homocysteine (SAH). Briefly, enzyme and substrates were incubated together along with the compound prior to quenching the reaction with 1% formic acid containing an internal standard of d4-SAH at 0.03 µg/mL. Unless stated otherwise, the experimental conditions used were the same as previously reported.^11^ The injection volume was 33 µL with 11 seconds (0.18 mins) per sample.

The experiment was modified later to use an additional internal standard (peptide IAYLKK*AT). Results were expressed as a normalised response; a ratio of the detected analyte, SAH, to the internal standard (either d4-SAH or IAYLKK^*^AT). A percent effect was calculated by comparing the compound well results to the 100% effect or 0% effect of the enzyme in the control wells.

### MTase-glo assay

The MTase-glo assay (Promega Corporation, Madison, WI, Cat. number V7601) was used as an orthogonal method of measuring nsp14 activity for confirming hits from the screen. Buffer conditions and proteins and substrate concentrations were kept the same as in the previously reported Rapid Fire assay.^11^ The assay was carried out in white flat bottom assay plates and was stopped by the addition of MTase-Glo reagent (1x) at the end of the incubation time. Plates were then incubated in the dark for 30 minutes prior to the addition of the MTase-Glo Detection Reagent. The signal was allowed to develop for 30 minutes in the dark and plates were then read using a PHERAstar FS instrument to detect luminescence.

### Pipeline pilot analysis

An automated Pipeline Pilot script was used to convert the internal standard and analyte data in the normalised response, as well as outputting the data in a standard 384 plate format. This serves to confirm successful sipping, successful detection of analyte by the RapidFire and smooth any well-to-well variability.

## Results and discussion

With the reported RF MS nsp14 assay^11^ in hand, we have screened libraries of compounds in a high-throughput manner for nsp14 inhibitors. Typically, the deuterated product d4-SAH is used as an internal standard in the experiment as it possesses similar properties in the mass spectrometer to that of the analyte product SAH. The expected behaviour is that the SAH signal will decrease with increasing inhibitor concentration, with the internal standard d4-SAH remaining at a constant level.

The SAH signal is conveniently normalised to the internal standard d4-SAH signal as a ratio, to gauge the degree of nsp14 inhibition. During the process of validating hits from this nsp14 screen, we observed a small set of compounds showing aberrantly high values for the internal standard d4-SAH when viewing live data collection via the Agilent MassHunter Workstation Data Acquisition Software, **Figure 1**. An automated analysis of these compounds showed well-behaved sigmoidal curves (Figure 2A). However, when these compounds were subsequently tested in an orthogonal activity assay (MTase-Glo) they were found to be inactive or had a significantly reduced activity (Figure 2D).

**Figure 1.**
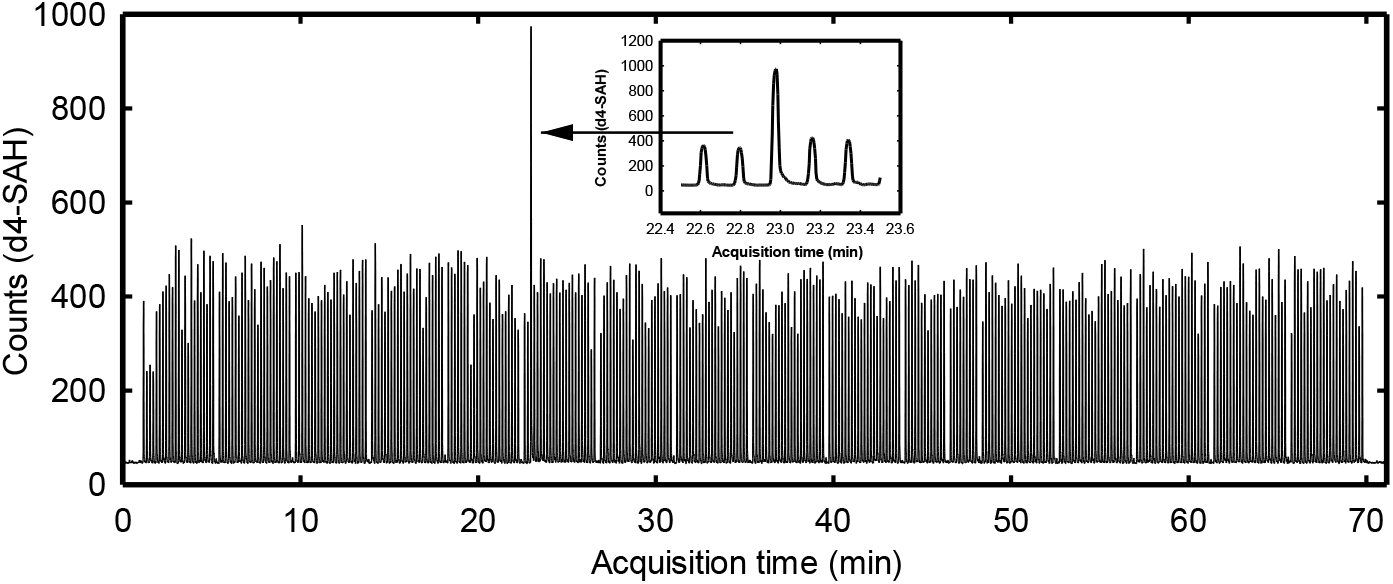
Example trace of high internal standard. Multiple reaction monitoring (MRM) chromatogram of d4-SAH from 384-well NSP14 assay with anomalous peak. Inset: Detail showing highlighted peak.

**Figure2.**
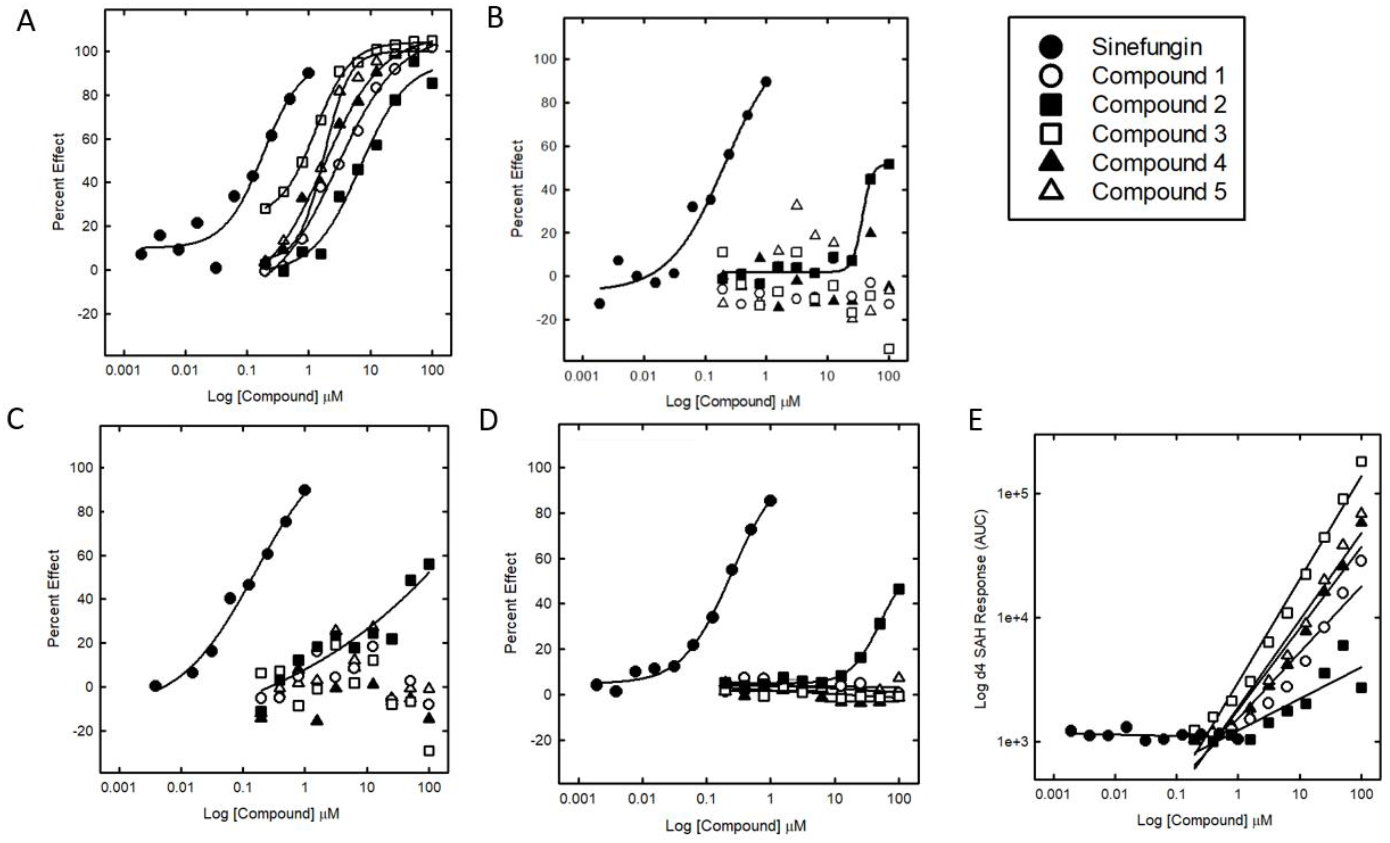
Dose response curves of the 5 test compounds and the Sinefungin control tested in singlicate. All percent effects (PE) are calculated with reference to 100% effect (0 nM nsp14+DMSO) and 0% effect wells (5 nM nsp14+DMSO)

To further understand the mechanism of these RF false-positive compounds, we re-examined the associated RF data in more detail. Inspection of the RF data revealed that the false-positive samples showed an unexpected dose-dependent increase in detected d4-SAH, rather than constant, and no dose-dependent decrease in SAH, or a significantly reduced decrease. When the SAH signal is normalised to the increasing d4-SAH signal, this gave the appearance of inhibition due to the compound-dependent decreasing SAH:d4-SAH ratio. If the analysis ignores the internal standard, and instead the percent effect is calculated using the effect on the SAH signal alone, these compounds showed either no apparent, or significantly reduced inhibition (see Figure 2b). Examples of compound structures which displayed this interference and their parent and predicted daughter ion m/z values are shown in **Table 1**.

**Table 1.**
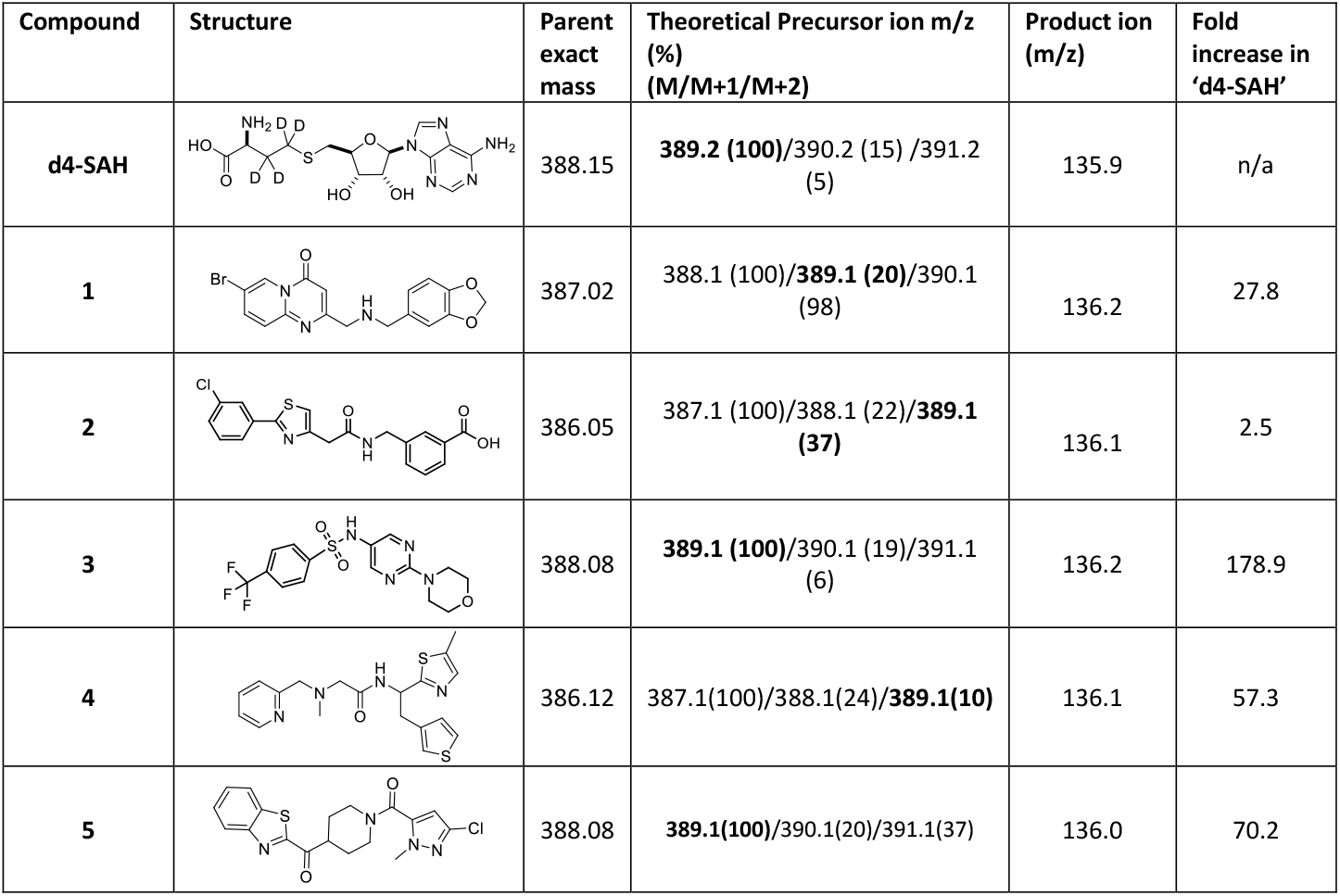
Compound structures and corresponding theoretical precursor ion with isotopic distribution and related product ion found in the d4-SAH measurement condition at precursor m/z 389.2(Figure S1-S5). Fold increase in d4-SAH is calculated using the mean of the d4-SAH in the control wells vs the detected d4-SAH inthe top concentration of the compound wells.

| Compound | Structure | Parent exact mass | Theoretical Precursor ion m/z (%) (M/M+1/M+2) | Product ion (m/z) | Fold increase in 'd4-SAH' |
| --- | --- | --- | --- | --- | --- |
| **d4-SAH** |  | 388.15 | **389.2 (100)**/390.2 (15) /391.2 (5) | 135.9 | n/a |
| **1** |  | 387.02 | 388.1 (100)/**389.1 (20)**/390.1 (98) | 136.2 | 27.8 |
| **2** |  | 386.05 | 387.1 (100)/388.1 (22)/**389.1 (37)** | 136.1 | 2.5 |
| **3** |  | 388.08 | **389.1 (100)**/390.1 (19)/391.1 (6) | 136.2 | 178.9 |
| **4** |  | 386.12 | 387.1(100)/388.1(24)/**389.1(10)** | 136.1 | 57.3 |
| **5** |  | 388.08 | **389.1(100)**/390.1(20)/391.1(37) | 136.0 | 70.2 |

Strikingly, all the false-positive hits have not only a precursor ion very close in mass to the internal standard d4-SAH but also a product ion that also closely matches the product ion of d4-SAH. Retesting of these compounds with an alternative internal standard (peptide IAYLKK*AT) showed that the false-positive compounds were indeed either completely inactive or had significantly reduced activity as expected. It seems therefore that the observed original levels of inhibition with d4-SAH as an internal standard was an artefact, caused by interference of the test compound and the detection of d4-SAH (see Figure 2). It is worth noting that compound 2 displayed some inhibition against nsp14 once the interference mechanism was eliminated. Although this compound did display a dose dependent increase in apparent d4-SAH detected by the RapidFire (Figure 2E), this was lower than that of the other compounds, and it still shows detectable inhibition when tested using the alternative internal standard (Figure 2C) and the MTase Glo orthogonal assay (Figure 2D). This is another example of the impact that certain types of interference can have as, although not completely a false positive, the true activity of the molecule is masked by its interference. This could potentially impact the design of more potent analogues and understanding of its SAR (structure activity relationship), as well as introducing the risk that any new molecules from this starting point would only be optimising towards its interference.

A. PE using SAH/d4-SAH normalised signal
B. PE using SAH signal (ignoring d4-SAH internal standard)
C. PE using SAH/peptide IAYLKK*AT normalised signal
D. PE in MTase Glo assay
E. d4-SAH signal

It may not necessarily be easy to predict if a compound is likely to interfere in this way based upon the compounds structure. However, if the internal standard used is close in molecular weight to that of the test compound, then it is advisable to consider this interference mechanism and treat any observed inhibition with caution. An orthogonal assay or rescreening the hits with a different internal standard will help to distinguish true versus false positives.

Although our initial identification was revealed by manual inspection of the raw data, we have now implemented a Pipeline Pilot script that can be used to identify compound wells with an unusually high internal standard value.

## Conclusion

In this work we have reported a previously unpublished potential mode of assay interference when using RapidFire MS technology and offer suggested mitigating strategies to either identify or avoid this. The ideal internal standard is one that is closely related to the enzyme product, which in the case of a methyltransferase and SAH, but this has a mass that is close to typical HTS screening compounds. In such cases, either implementing an automated script to monitor the internal standard signal or flagging hit compounds with masses close to the internal standard is recommended to be included as an additional quality control layer. If a second internal standard with a molecular weight beyond the range of HTS compounds can be practically included in the experiment, that can help circumvent compound interference issues. In conclusion, the RF-MS screening method remains a powerful technique that is anticipated to be less susceptible than other methods to false-positives, however caution is advised when hit compounds are close in molecular weight to the internal standard.

## Acknowledgements

This work was supported by the Wellcome Trust [203134/Z/16/Z] and the Bill & Melinda Gates Foundation [INV-016131].We would also like to acknowledge the assistance of the DDU Compound Management Team (Alex Cookson, Kirstie Cookson, Fraser Hughes, Steve Bell) and the DDU Data Management Team (Stephen Thompson, Kashish Sharma, Edan Gardner) for their assistance, and Alana Pineiro (DDU) for her abstract.

## Notes

### Competing Interest Statement

The authors have declared no competing interest.

