## Supplementary Information for "Identification of a novel false-positive mechanism in RapidFire mass spectrometry and application in drug discovery"

† Joint first author    * Corresponding author

Contents: Figures S1-S5

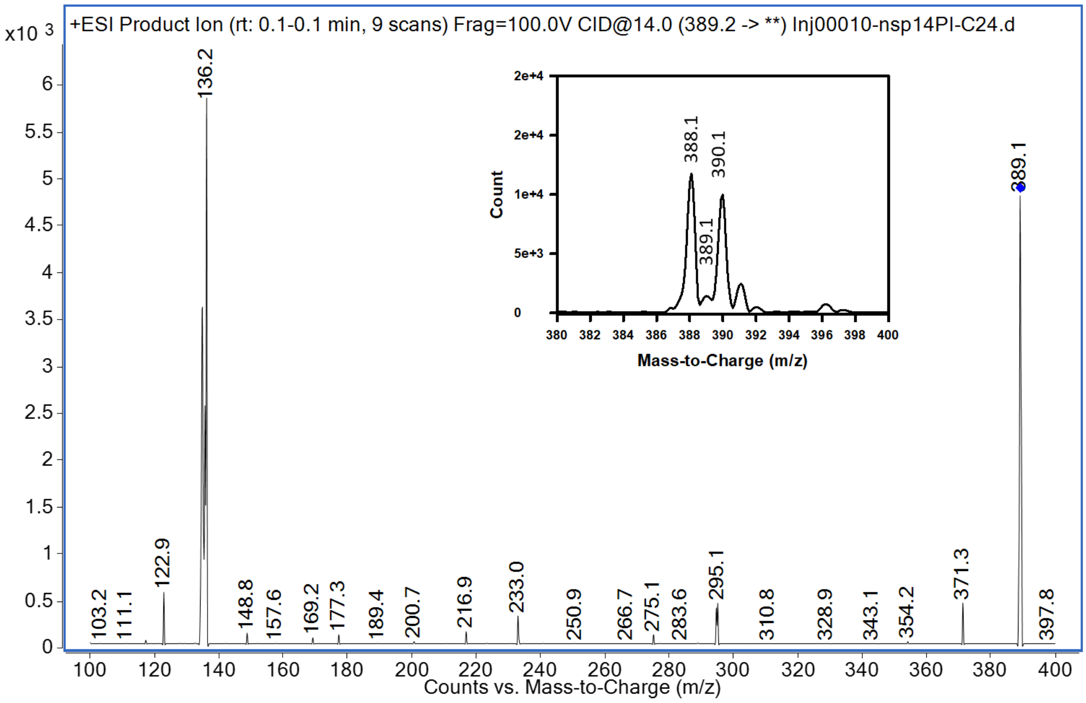


Figure S1: Compound 1 product ion scan of m/z 389.2, scans were acquired from m/z 100-400. The inset shows the full scan (m/z 380-400) with relative abundances of M (m/z 388.1), M+1 (m/z 389.1), and +2 (m/z 390.1) isotopes.

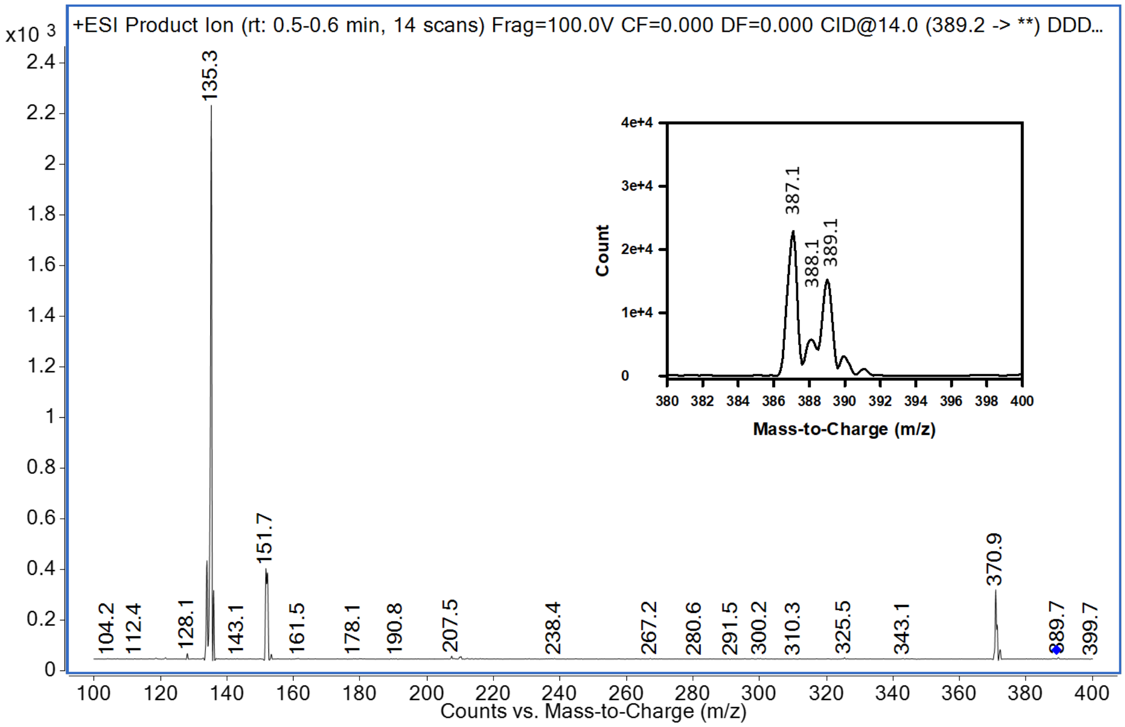


Figure S2: Compound 2 product ion scan of m/z 389.2, scans were acquired from m/z 100-400. The inset shows the full scan (m/z 380-400) with relative abundances of M (m/z 387.1), M+1 (m/z 388.1), and +2 (m/z 389.1) isotopes.

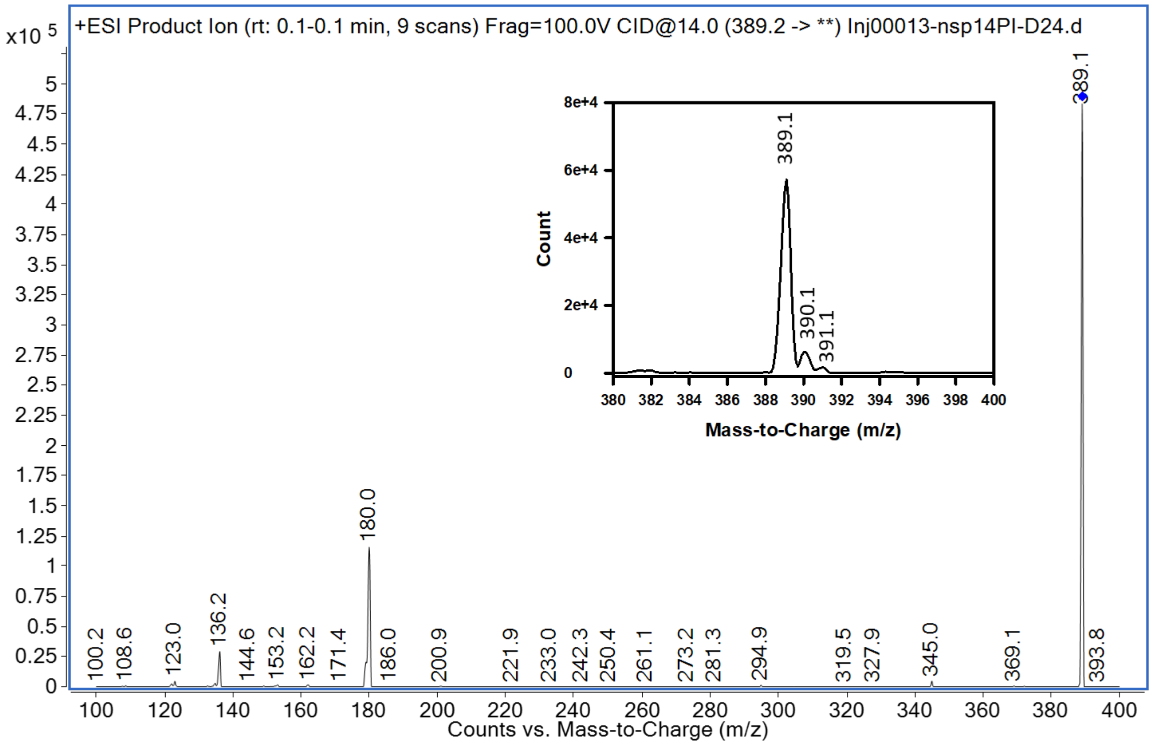


Figure S3: Compound 3 product ion scan of m/z 389.2, scans were acquired from m/z 100-400. The inset shows the full scan (m/z 380-400) with relative abundances of M (m/z 389.1), M+1 (m/z 390.1), and +2 (m/z 391.1) isotopes.

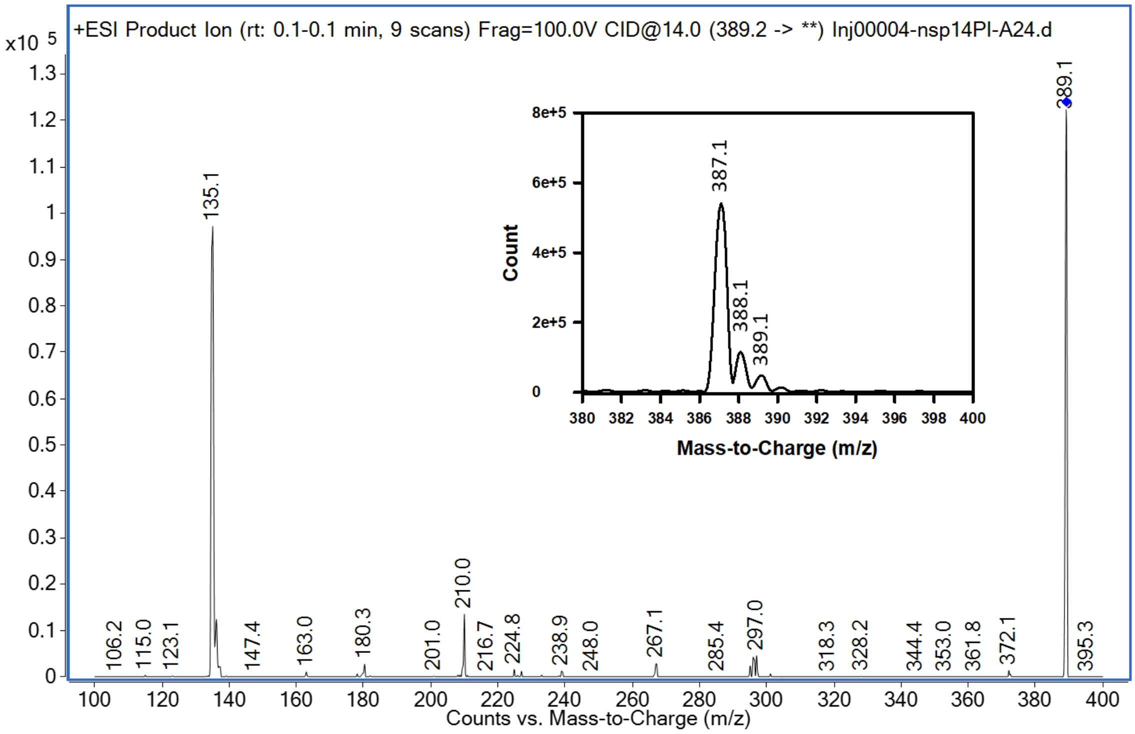


Figure S4: Compound 4 product ion scan of m/z 389.2, scans were acquired from m/z 100-400. The inset shows the full scan (m/z 380-400) with relative abundances of M (m/z 387.1), M+1 (m/z 388.1), and +2 (m/z 389.1) isotopes.

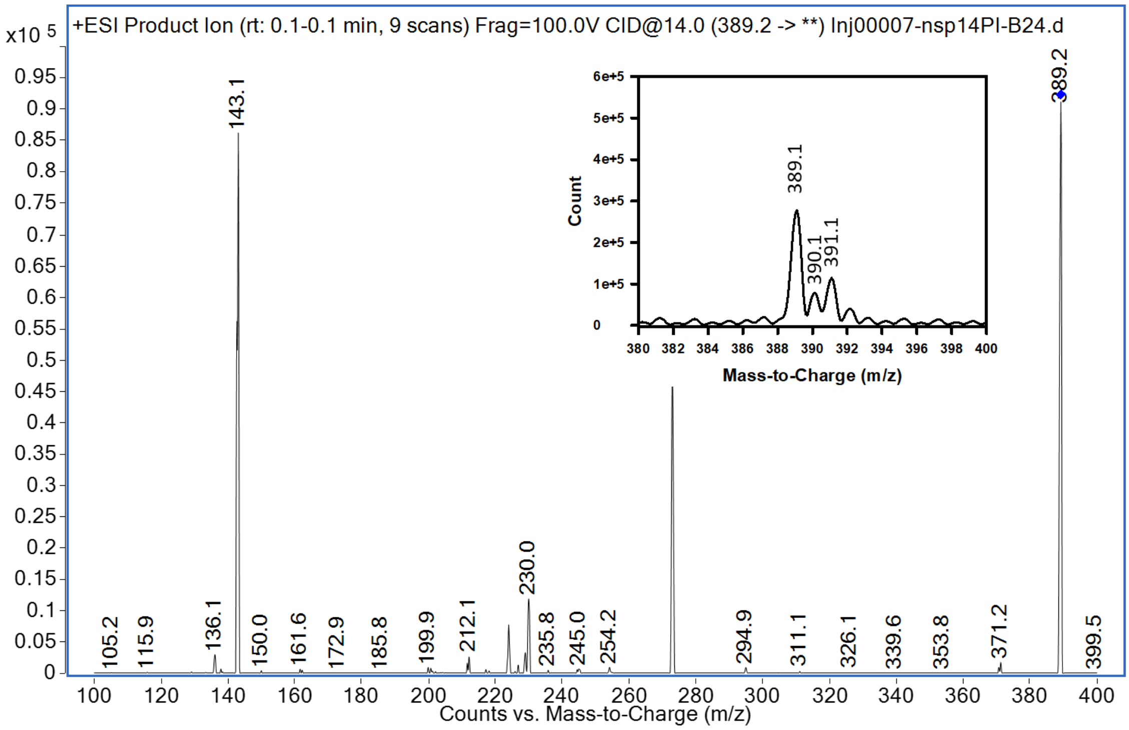


Figure S5: Compound 5 product ion scan of m/z 389.2, scans were acquired from m/z 100-400. The inset shows the full scan (m/z 380-400) with relative abundances of M (m/z 389.1), M+1 (m/z 390.1), and +2 (m/z 391.1) isotopes.
